# Assortative Mating, Autistic Traits, Empathizing, and Systemizing

**DOI:** 10.1101/2020.10.28.358895

**Authors:** Gareth Richards, Simon Baron-Cohen, Holly Stokes, Varun Warrier, Ben Mellor, Ellie Winspear, Jessica Davies, Laura Gee, John Galvin

**Author notes:** Corresponding author address: School of Psychology, Newcastle University, 2.27 Ridley Building 1, Queen Victoria Road, Newcastle upon Tyne, UK.

## Abstract

It has been suggested that the children of parents with particular interests and aptitude for understanding systems via input-operation-output rules (i.e. systemizing) are at increased likelihood of developing autism. Furthermore, assortative mating (i.e. a non-random pattern in which individuals are more likely to pair with others who are similar to themselves) is hypothesised to occur in relation to systemizing, and so romantic couples may be more similar on this variable than chance would dictate. However, no published study has yet tested this hypothesis. We therefore examined intra-couple correlations for a measure of autistic traits (Autism Spectrum Quotient [AQ]), self-report measures of empathizing (Empathy Quotient [EQ]), and systemizing (Systemizing Quotient-Revised [SQ-R]), as well as the Reading the Mind in the Eyes Test (RMET) and Embedded Figures Task (EFT). We observed positive intra-couple correlations of small-to-medium magnitude for all measures except EQ. Further analyses suggest that these effects are attributable to people pairing with those who are more similar to themselves than chance (initial assortment) rather than becoming more alike over the course of a relationship (convergence), and to seeking out self-resembling partners (active assortment) rather than pairing in this manner due to social stratification increasing the likelihood of similar people meeting in the first place (social homogamy). Additionally, we found that the difference in scores for the AQ, SQ-R, RMET and EFT of actual couples were smaller (i.e. more similar) than the average difference scores calculated from all other possible male-female pairings within the dataset. The current findings therefore provide clear evidence in support of the assortative mating theory of autism.

## 1. Introduction

Autism spectrum conditions are characterised by unusually routine behaviours, narrow interests, sensory differences, and social and communicative difficulties (American Psychiatric Association, 2013). There is a marked sex difference in autism diagnosis, with approximately four males being diagnosed per every one female (Baio et al., 2018; Fombonne, 2009), an effect that may (at least in part) be explainable in terms of biological factors such as atypical foetal sex hormone exposure (Auyeung et al., 2009; Baron-Cohen et al., 2015, 2019) and social stereotyping processes (Bargiela et al., 2016; Geelhand et al., 2019; Whitlock et al., 2020). The prevalence of autism is approximately 1-2% (Baio et al., 2018), which is a marked increase from Lotter’s (1966) early estimate of 0.045%. This increase in prevalence may be explained by better recognition of autistic symptoms, growth of relevant services, diagnostic upgrading (e.g. providing a diagnosis when uncertain and/or to ensure school support or access to services), and broadening of diagnostic criteria (Arvidsson et al., 2018; Dawson, 2013; Fombonne, 2009, 2018; Roelfsema et al., 2012). However, it remains unclear whether such processes can explain the rise in prevalence in its entirety or whether the actual incidence of autism has also increased in recent years (Fombonne, 2009).

With excellent attention to detail, and the ability to remain focussed on repetitive tasks, autistic people often have an aptitude for working in Science, Technology, Engineering, and Mathematics (STEM) industries (Baron-Cohen et al., 1997, 2007; Baron-Cohen, Wheelwright, Skinner, et al., 2001; Ruzich et al., 2015; Wei et al., 2013). This effect also appears to extend to the broader autism phenotype (BAP) - those people who display higher than average levels of autistic traits but do not warrant a clinical diagnosis (Baron-Cohen, Wheelwright, Skinner, et al., 2001; Hoekstra et al., 2008; Pisula et al., 2013; Wakabayashi et al., 2006). Furthermore, it has been suggested that autism might be subject to positive assortative mating (i.e. people who measure highly in autistic traits may be more likely than chance to have children together; Baron-Cohen, 2006b, 2006a, 2007). However, there is more than one process by which this could operate. For instance, it may be that individuals with similar levels of autistic traits consciously or unconsciously seek each other out as romantic partners (active assortment) or that individuals with similar levels of autistic traits are more likely than chance to share other characteristics, such as a working environment, which may lead to an increased likelihood of a relationship starting (social homogamy); in addition, individuals may begin relationships with others who are more similar to themselves than expected by chance as regards autistic traits (initial assortment) or become more similar to their partner over the course of their relationship (convergence) (Kardum et al., 2017; Luo, 2017).

Evidence for autism being subject to assortative mating comes from the observation that a person diagnosed with autism is 10-12 times more likely to marry or have a child with another autistic person than is someone without such a diagnosis (Nordsletten et al., 2016). It should, however, be noted that due to matching five controls to every case, the partner correlation (*r* = 0.47) observed by Nordsletten et al. (2016) has been estimated to reflect a smaller effect (~*r* = 0.28) in the general population (Peyrot et al., 2016). Furthermore, although a study utilising genome wide association study (GWAS) data (Yengo et al., 2018) lacked the statistical power required to detect a significant genotypic correlation, a more recent study by Connolly et al. (2019) reported greater genetic similarity than expected by chance in the parents of autistic children. Additionally, several studies have reported positive intra-couple correlations for phenotypic autistic traits measures (Connolly et al., 2019; Constantino & Todd, 2005; Duvekot et al., 2016; Hoekstra et al., 2010; Lyall et al., 2014; Schwichtenberg et al., 2010; Virkud et al., 2009), although others have reported null (Hoekstra et al., 2007; Losh et al., 2008; Pollmann et al., 2010; Van Steijn et al., 2012) or ambiguous findings (Seidman et al., 2012). Meta-analysis of the available literature (k=15, n=5,770 [including the current study]) shows a small but highly statistically significant positive correlation, *r* = 0.200 (95% CI = 0.109, 0.289), *p* < 0.0001 (see **Appendix**). Interestingly there appears to be no published research examining intra-couple correlations for measures of empathizing (Baron-Cohen & Wheelwright, 2004) and systemizing (Wheelwright et al., 2006), despite it having been hypothesised that autism may result from the pairing of two high systemizers (Baron-Cohen, 2007).

The current study aims to increase our understanding of the processes that might underpin assortative mating as it relates to autism. More specifically, we examined intra-couple correlations for quantitative self-report measures of autistic traits (Autism Spectrum Quotient [AQ]; Baron-Cohen, Wheelwright, Skinner, et al., 2001), empathizing (Empathy Quotient [EQ]; Baron-Cohen & Wheelwright, 2004), and systemizing (Systemizing Quotient-Reived [SQ-R]; Wheelwright et al., 2006), as well as the standardised difference between empathizing and systemizing (D scores). Additionally, we examined behavioural measures that broadly map onto empathizing (Reading the Mind in the Eyes Test [RMET]; Baron-Cohen, Wheelwright, Hill, et al., 2001) and systemizing (Embedded Figures Test [EFT]; Witkin et al., 1971). We predicted that: (1) sex differences would be observed for each of these measures (M>F for AQ, SQ-R, D score, and EFT; F>M for EQ and RMET), (2) variables associated with social homogamy (age, educational attainment, and STEM status) would correlate positively within couples, (3) autism-related measures would be positively correlated within couples; we also predicted (4) that intra-couple correlations for autism-related variables would reflect initial assortment rather than convergence, (5) that intra-couple similarity for autism-related variables might reflect social homogamy for age and attainment or active assortment (no directional prediction was made here), and (6) that intra-couple correlations for autism-related variables would be stronger in couples for whom both partners worked/studied in STEM.

## 2. Method

### 2.1 A priori power analysis

We conducted an *a priori* power analysis using G*Power 3.1 (Faul et al., 2007, 2009) to determine the sample size. Assuming a medium effect size (*r* = 0.30) for intra-couple correlations on personality variables (e.g. Kardum et al., 2017) and 80% power, this analysis determined that a sample size of n=67 couples would be required to observe a statistically significant effect (*p* < 0.05) with a one-tailed Pearson’s correlation test.

### 2.2 Apparatus/Materials

#### 2.2.1 Demographics

Participants firstly reported their sex (‘Male’, ‘Female’, ‘Prefer not to say’), ‘Other (please specify below)’, age (years), and ethnicity (‘White’, ‘Mixed / multiple ethnic groups’, ‘Asian / Asian British’, ‘Black / African / Caribbean / Black British’, ‘Other ethnic group’). They were then asked questions regarding their relationship, specifically their cohabiting status (‘Living with partner’, ‘Not living with partner’), length of relationship (years and months), and marital status (‘Not married’, ‘Engaged to be married’, ‘Married’). They were also asked to confirm their educational level (‘No qualifications’, ‘Completed GCSE level (or equivalent)’, ‘Completed A level, Access Course (or equivalent)’, ‘Bachelor’s Degree’, ‘Master’s Degree’, ‘Doctorate Degree’, ‘Other, please specify)’), current student status (‘Yes’, ‘No’; if ‘Yes’, then area and year of study were also recorded), whether they were employed (‘Yes’, No’; if ‘Yes’, then place of work and job role were also recorded), whether they had an autism diagnosis (‘Yes’, ‘No’) or suspected they were autistic (‘Yes’, ‘No’). See **Table 1** for descriptive statistics relating to the sample’s demographics.

#### 2.2.2 Autistic traits and related measures

We used the 50-item self-report Autism Spectrum Quotient (AQ; Baron-Cohen et al., 2001) to measure autistic traits. For each item, participants are asked to specify to what extent they consider a statement to relate to them (response options: Strongly agree, Slightly agree, Slightly disagree, Strongly disagree). Half of the items are reverse-coded, and one point is given for each response (either slight or strong) validating an autistic trait. The sum of all 50 items (possible range = 0-50) is calculated as an indicator of one’s level of autistic traits (higher scores indicate more autistic traits). Cronbach’s alpha was considered satisfactory (i.e. > 0.70; Bland & Altman, 1997; see also Tavakol & Dennick, 2011) in the current study (α = 0.823).

We measured self-reported empathizing via the 40-item Empathy Quotient (EQ; Baron-Cohen & Wheelwright, 2004). Although the response options are the same as those of the AQ, in this case, participants are assigned 1 point for each response that slightly endorses an empathic tendency and 2 points for a response that strongly endorses an empathizing tendency. Approximately half of the items are reverse-scored, and the possible range of scores is 0-60 (higher scores indicate higher empathizing). Internal consistent for this measure was satisfactory (α = 0.923). Self-reported systemizing was measured via the 75-item Systemizing Quotient-Revised (SQ-R; Wheelwright et al., 2006). As with the EQ, 1 point is assigned for each response slightly endorsing a systemizing tendency and 2 points are assigned for each response strongly endorsing a systemizing tendency; scores can range from 0 to 150, and higher scores indicate higher systemizing. The SQ-R showed satisfactory internal consistency in the current study (α = 0.911). In addition to examining EQ and SQ-R scores, we also standardised these scores and calculated the difference as: D = S – E. D scores provide an indication of one’s cognitive style, with positive scores indicating relatively strong systemizing compared to empathizing.

In addition to the questionnaires, two behavioural measures were administered. Firstly, the Reading the Mind in the Eyes Test (Baron-Cohen, Wheelwright, Hill, Raste, & Plumb, 2001) was used to provide an indication of participants’ ability to correctly infer mental states in others, a skill that broadly maps onto empathizing. For this task, a picture of the eye region is shown along with four adjectives, one of which correctly describes the emotion portrayed; a practice trial is completed before the 36 items that comprise the measure. Internal consistency was satisfactory (α = 0.709). In addition, we used the Embedded Figures Test (Witkin et al., 1971) as a behavioural task that taps into abilities prerequisite to systemizing. For this measure, participants are shown a rectangular stimulus consisting of horizontal, vertical, and diagonal lines, within which they are tasked with identifying a particular shape (i.e. the embedded figure). A practice trial is conducted prior to the 12 trials that comprise the measure. Participants are timed on each trial by a Research Assistant using a stopwatch, and the time taken to identify the correct shape is recorded (if the participant does not identify the correct shape within 180 seconds, the task proceeds to the next trial). The mean score across all 12 trials is computed as the variable of interest. The internal consistency for this measure was satisfactory (α = 0.787).

### 2.3 Design and Procedure

The current study utilised a correlational design. Participants were invited to attend a lab session in which they and their partner would independently complete several measures relating to autism. Each couple was offered a £10.00 Amazon voucher as an incentive to participate. The study protocol, hypotheses, and analysis plan were pre-registered on the Open Science Framework (OSF) prior to data analysis (osf.io/6jg8p). However, due to restrictions imposed by the COVID-19 pandemic, a minority of participants completed the study via an online survey hosted by Qualtrics. Digit ratio (2D:4D) was also measured for those participants that attended the lab, and results from that aspect of the study have been published elsewhere (Richards, Baron-Cohen, van Steen, & Galvin, 2020).

### 2.4 Statistical analyses

We first tested for sex differences in the autism-related variables by using independent samples *t* tests. We then examined intra-couple associations for age (Pearson’s correlation), educational attainment (Spearman’s correlation) and STEM study/occupational status (Chi-square test) before using Pearson’s correlations to determine the strength and direction of intra-couple correlation for each of the autism-related variables. Next, we compared the sex-standardised difference scores for autism-related variables for actual couples with those calculated by pairing each male in the dataset with each female other than his partner. This analysis was used to determine whether actual couples’ scores for these variables were more similar than expected assuming a pattern of random mating. We then used Pearson’s correlations to determine the level of association between length of relationship and the within-couple difference scores for autism-related variables. The assumption of this analysis was that a significant negative correlation would imply that couples’ scores become more similar over time and so indicate convergence effects rather than initial assortment. We then performed similar analyses in relation to age and educational attainment under the assumption that a significant positive correlation could indicate that couple similarity for autism-related variables is explainable in terms of social homogamy rather than active assortment. Finally, we used independent samples *t* tests to determine whether couples for whom both members worked/studied in STEM areas were more similar in terms of autism-related variables than couples for whom only one or neither member worked/studied in STEM.

We considered *p* < 0.050 (two-tailed) to suggest statistical significance and interpret effect sizes in accordance with Cohen (1988). Data analyses were conducted in R version 1.3.1073.

## 3. Results

### 3.1 Sex differences

As predicted, males scored higher than their female partners on AQ, SQ-R, and D score, and achieved faster times on the EFT; females had higher scores on the EQ and RMET, although the latter effect was not statistically significant (**Table 2**).

**Table 2.**
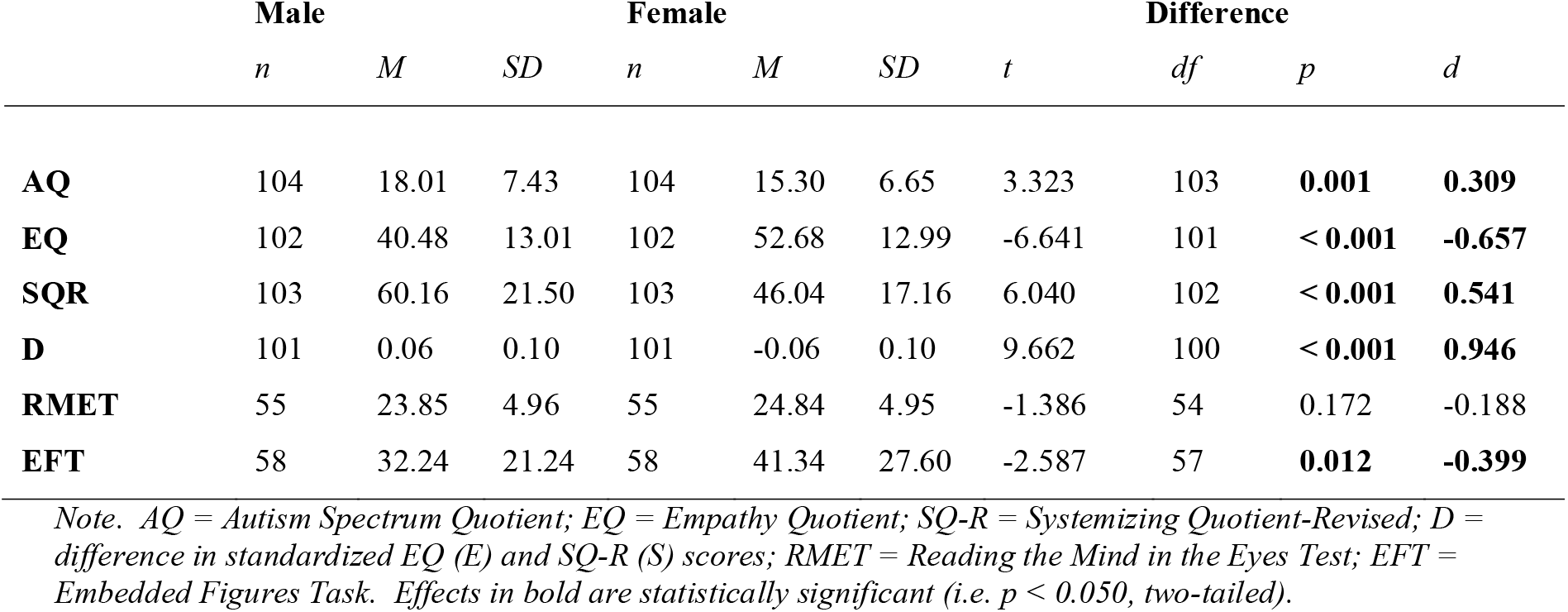
Descriptive statistics and sex differences for autism-related variables.

**Table 2.** Comparison of actual couple similarity for autism-related variables with the average difference scores calculated by pairing each participant all other potential opposite-sex partners in the dataset

|  | Actual couple difference |  | Average difference of all<br>other pairings |  | Difference |  |  |  |
| --- | --- | --- | --- | --- | --- | --- | --- | --- |
|  | *M* | *SD* | *M* | *SD* | *t* | *df* | *p* | *d* |
| **AQ** | 0.891 | 0.770 | 1.118 | 0.433 | -3.227 | 103 | **0.002** | **-0.250** |
| **EQ** | 1.109 | 0.861 | 1.111 | 0.384 | -0.025 | 101 | 0.980 | -0.002 |
| **SQR** | 0.928 | 0.779 | 1.128 | 0.410 | -2.761 | 102 | **0.007** | **-0.211** |
| **D** | 0.143 | 0.109 | 0.156 | 0.070 | -1.573 | 100 | 0.119 | -0.094 |
| **RMET** | 0.837 | 0.634 | 1.101 | 0.408 | -2.817 | 54 | **0.007** | **-0.393** |
| **EFT** | 0.788 | 0.722 | 1.071 | 0.520 | -2.847 | 57 | **0.006** | **-0.365** |

*Note.* *AQ* = Autism Spectrum Quotient; *EQ* = Empathy Quotient; *SQ-R* = Systemizing Quotient-Revised; *D* = difference in standardized *EQ* (*E*) and *SQ-R* (*S*) scores; *RMET* = Reading the Mind in the Eyes Test; *EFT* = Embedded Figures Task. Effects in bold are statistically significant (i.e. $p < 0.050$ , two-tailed).

### 3.2 Intra-couple correlations for social homogamy variables

Partners’ ages were very strongly positively correlated, *r*(103) = 0.964, *p* < 0.001. A Spearman’s correlation also demonstrated that partners’ level of educational attainment was positively correlated, *r*_s_(103) = 0.399, *p* < 0.001, and a Chi-square test showed that those studying/working in STEM were more likely than chance to have a partner who was also working/studying in STEM, χ^2^ (1, 87) = 11.481, *p* < 0.001, φ = −0.39.

### 3.3 Intra-couple correlations for autism-related variables

Positive intra-couple correlations were observed for AQ, *r*(102) = 0.305, *p* = 0.002, SQ-R, *r*(101) = 0.263, *p* = 0.007, D score, *r*(99) = 0.195, *p* = 0.051, RMET, *r*(53) = 0.438, *p* < 0.001, and EFT, *r*(56) = 0.423, *p* < 0.001, though not for EQ, *r*(100) = −0.018, *p* = 0.860 (**Figure 1**).

**Figure 1.**
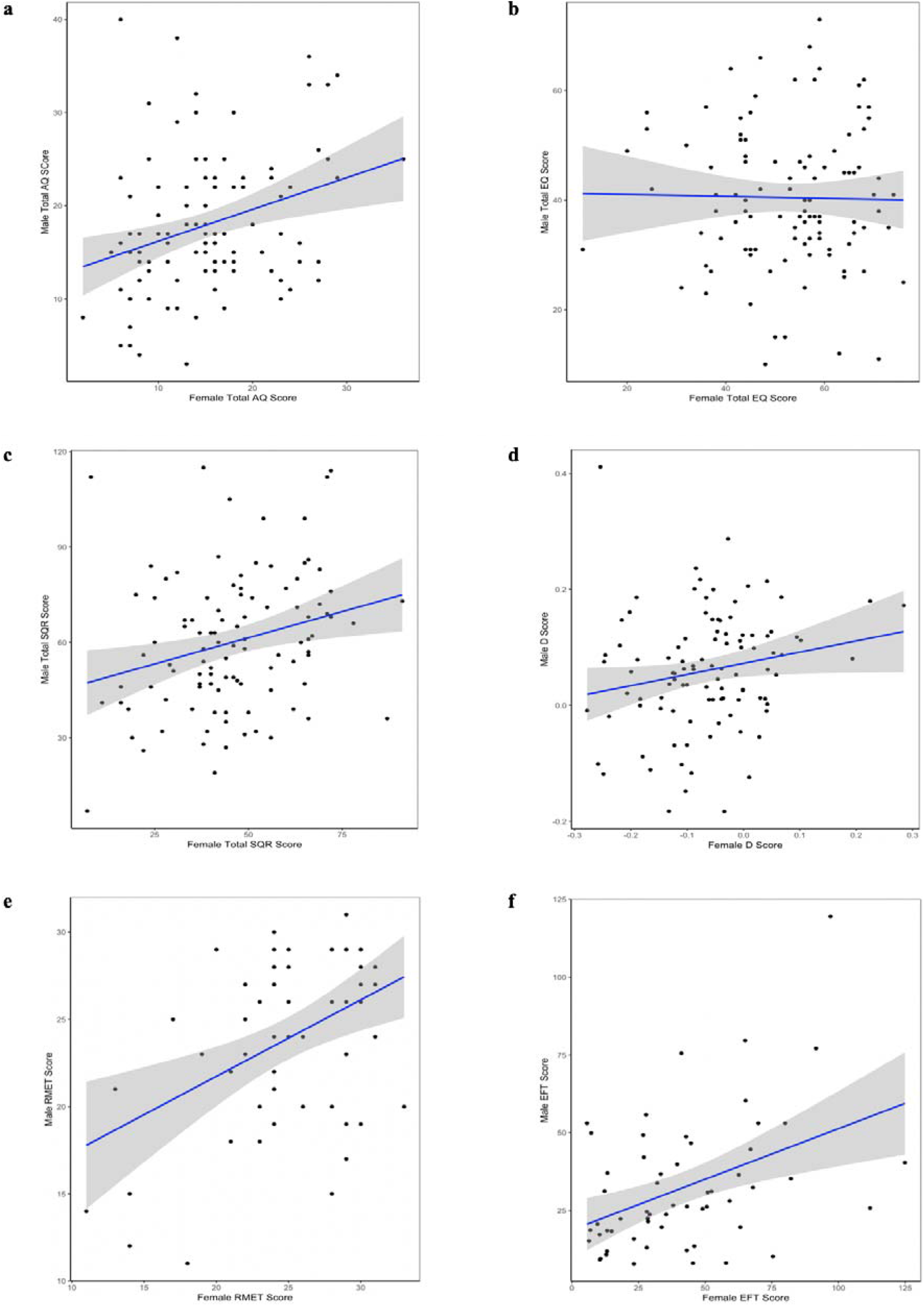
Scatterplots with regression line and 95% confidence intervals demonstrating the level of intra-couple correlation observed for AQ (a), EQ (b), SQ-R (c), D score (d), RMET (e), and EFT (f).

### 3.4 Couple similarity for autism-related variables

To determine whether actual couples within our dataset were more similar to each other on autism-related variables, we first calculated (unsigned) difference scores for each of the relevant variable as male score – female score. Next we calculated the difference scores between any given male and all females that were not his partner and took the average. Paired-samples *t* tests determined that the AQ, SQ-R, RMET and EFT difference scores for actual couples were smaller (i.e. more similar) than those calculated from random pairings, and that the observed differences were small in size; no such effects were observed for EQ and D scores (**Table 2**).

### 3.5 Initial assortment vs. convergence

To investigate whether the intra-couple correlations for autism-related variables were explainable by initial assortment or convenience, we correlated the standardised within-couple difference scores for autism-related variables with length of relationship. Essentially, if length or relationship is correlated with the difference score, it suggests that partners’ become more similar (negative correlation) or more dissimilar (positive correlation) over the course of their relationship, and so provides evidence against there being initial assortment (Kardum et al., 2017). As both males and females reported the length of their relationships, we correlated these two sets of scores to check for similarity (*r*[102] = 0.999, *p* < 0.001) before averaging the two measures for use in further analyses. The averaged measure showed a wide range of relationship lengths (from 1 month to 60 years, *M* = 9.365, *SD* = 11.033).

Relationship length was not correlated with the within-couple differences in AQ, *r*(102) = −0.051, *p* = 0.610, EQ, *r*(100) = −0.009, *p* = 0.929, SQ-R, *r*(101) = −0.012, *p* = 0.902, D, *r*(99) = 0.079, *p* = 0.430, RMET, *r*(53) = 0.153, *p* = 0.265, or EFT, *r*(56) = 0.079, *p* = 0.556. These findings therefore indicate that intra-couple correlations for autism-related variables are attributable to initial assortment rather than convergence effects.

### 3.6 Active assortment vs. social homogamy

To investigate whether intra-couple correlations for autism-related variables may be better explained by active assortment or social homogamy, we first correlated the within-couple standardised difference scores for the autism-related variables with the couples’ absolute difference in age. The idea here is that if couples that are dissimilar in age are also more dissimilar for autism-related variables, the intra-couple correlation may be explained by social homogamy effects. We observed no correlation between absolute (i.e. unsigned) age difference and within-couple differences for AQ, *r*(102) = −0.016, *p* = 0.870, SQ-R, *r*(101) = 0.009, *p* = 0.928, D, *r*(99) = −0.125, *p* = 0.212; RMET, *r*(53) = 0.225, *p* = 0.099, or EFT, *r*(56) = 0.022, *p* = 0.870. However, there was a significant correlation for EQ, *r*(100) = −0.244, *p* = 0.014. This effect suggests that couples with disparate age-gaps may be more similar in empathizing than expected by chance. We next examined whether there were correlations between couple similarity for level of educational attainment and autism-related variables. Couple similarity for educational attainment was not associated with couple similarity for AQ, *r*_s_ (102) = −0.065, *p* = 0.514, EQ, *r*_s_ (100) = −0.048, *p* = 0.634, SQR, *r*_s_ (101) = 0.084, *p* = 0.400, D, *r*_s_ (99) = −0.040, *p* = 0.694, or EFT, *r*_s_ (56) = 0.166, *p* = 0.212; although couples who were more similar for educational attainment were also more similar than expected for RMET scores, the effect was not statistically significant, *r*_s_ (53) = 0.239, *p* = 0.078. Finally, we examined whether couples who work in STEM are more similar on autism-related traits than are couples for whom either one member or both members do not work in STEM. There were 35 males and 33 females who worked or studied in STEM areas, and 57 males and 61 females who did not study or work in STEM areas. We compared the 20 couples for whom both members were in STEM with the 67 couples for whom either one or neither member was in STEM. These groups of couples did not differ in regard to their level of similarity on the AQ, *t*(37.315) = −0.464, *p* = 0.646, EQ, *t*(45.323) = 0.304, *p* = 0.762, SQ-R, *t*(37.327) = −0.072, *p* = 0.943, D, *t*(33.19) = −0.003, *p* = 0.998, RMET, *t*(9.925) = 0.269, *p* = 0.793, or EFT, *t*(12.736) = −0.197, *p* = 0.847 (equal variances were not assumed in each case). The overall pattern of results observed here is consistent with active assortment, though some effects of social homogamy cannot be ruled out.

## 4. Discussion

The current study aimed to provide an empirical examination of the assortative mating theory of autism by determining if (and to what extent) traits associated with autism are correlated within heterosexual partners in the general population. The main finding was that quantitative measures of autistic traits (AQ) and systemizing (SQ-R), the standardised difference between empathizing and systemizing (D), the ability to read emotions in the eye region (RMET), and spatial skills (EFT) (but not empathizing; EQ) are all positively correlated within partners, and that these effects are better explained in terms of active assortment than social homogamy, and by initial assortment rather than convergence. Furthermore, when we compared within-couple standardised difference scores for these variables with standardised difference scores calculated as the average of all other possible heterosexual pairings within the dataset, we found that actual couples were more similar for AQ, SQ-R, RMET, and EFT than would be expected under the assumption of random mating.

A number of studies have previously examined intra-couple correlations for autistic trait variables, the majority of which have focused on very specific samples such as the parents of autistic children (Connolly et al., 2019; Lau et al., 2014; Losh et al., 2008; Lyall et al., 2014; Schwichtenberg et al., 2010; Seidman et al., 2012; Van Steijn et al., 2012; Virkud et al., 2009), parents of twins (Constantino & Todd, 2005; Hoekstra et al., 2010; Hoekstra et al., 2007), and parents of children with various mental health conditions (Duvekot et al., 2016). Although two of these have also included samples of parents of typically developing children (Lyall et al., 2014; Schwichtenberg et al., 2010) and another study reported on a sample of newlyweds (Pollmann et al., 2010), it is clear that the nature of these effects within the general population remains relatively unexplored. As quantitative autistic traits share a genetic architecture with autism in terms of a diagnostic construct, children of two parents who both have high levels of autistic traits can be predicted to receive a double dose in terms of polygenetic susceptibility to autism. This may have important implications as regards our understanding of geographical trends in autism prevalence. This point is made clearer by the finding that rates of autism can sometimes cluster (Mazumdar et al., 2010; van Meter et al., 2010), with children in Eindhoven (i.e. the ‘Silicon Valley of Europe’) being twice as likely to be autistic as children from two similar sized cities (Utrecht and Haarlem) in the Netherlands (Roelfsema et al., 2012). Furthermore, Peyrot et al. (2016) suggested that “current trends in assortative mating might lead to a considerable increase in the prevalence of rare disorders with high heritability”.

A novel aspect of the current study is that we demonstrated statistically significant positive intra-couple correlations (and increased within-couple similarity) for self-reported systemizing (SQ-R scores) as well as for a behavioural/cognitive skill that likely underpins systemizing ability (speed on the EFT). This is particularly important considering that systemizing shares a genetic architecture with autism (Warrier et al., 2019), and, of course, because it is the trait upon which assortative mating in relation to autism was initially hypothesised to act (Baron-Cohen, 2006a, 2006b, 2007). These findings may therefore be informative regarding why autism is associated with interest and aptitude for STEM (Baron-Cohen et al., 1997, 2007; Baron-Cohen, Wheelwright, Skinner, et al., 2001; Ruzich et al., 2015; Wei et al., 2013), and why children in geographical regions enriched with STEM industry are at elevated likelihood of developing the condition (Roelfsema et al., 2012).

It remains unclear exactly why autistic traits and systemizing are correlated within couples, though it may be that increased similarity for these variables improves the ‘flow’ of a relationship and therefore helps it survive and progress. The observation that these associations were not moderated by length of relationship or social homogamy variables suggests that they likely reflect initial and active assortment (i.e. people seek out similar individuals as partners and do not become more alike over the course of their relationship). Although Pollmann et al. (2010) reported that autistic traits were not associated with relationship satisfaction in wives, husbands with high levels of autistic traits had lower relationship satisfaction, an effect that was entirely mediated by low trust in and responsiveness to their partner, and low intimacy in the relationship. Perhaps somewhat counterintuitive though is the observation of Jobe and Williams White (2007) that autistic traits are actually positively correlated with relationship length, though this might be explained by those with high levels of autistic traits typically being resistant to change and so less likely to choose to end a relationship. However, although speculative, this resistance to change might also result in couples being less likely to converge on these measures over the course of their relationship, which provides further support for the presence of initial assortment.

It is relevant to note that similarity in people’s levels of autistic traits extends beyond romantic relationships, as it has also been observed in friendship dyads. Wainer, Block, Donnellan, and Ingersoll (2013) reported that autistic traits (as measured by the BAPQ) were significantly correlated between same-sex friendship pairs, and that this effect was present when examined for both self-report (*r* = 0.23) and informant-(i.e. partner) report measures (*r* = 0.16). Of particular relevance here is the finding that concordance on autism-related traits (specifically the ‘aloofness’ scale of the BAPQ) predicted increased relationship satisfaction in newly-formed college roommate dyads when measured at 9-10-week follow-up (Faso et al., 2016). Furthermore, autistic adults appear to be more comfortable during first interactions with other autistic (as opposed to neurotypical) adults (Morrison et al., 2020). Taken together, these findings imply that individuals with high levels of autistic traits find it easier to begin relationships with people who show concordance in this regard, and that such relationships are more likely to progress. This process may therefore also be implicated in the development of romantic relationships, with individuals more closely matched on autistic traits being more likely than discordant dyads to pursue and maintain them.

An important consideration in explaining the underpinnings of assortative mating is the possible role that online dating may play. In particular, evidence suggests that online dating relates to reduced assortative mating in regard to occupation and geographical proximity but increased couple similarity for educational level, age, and marital history (Lee, 2016). The Internet provides a novel environment in which to seek potential romantic partners. Like never before in our evolutionary history, it provides opportunities for individuals to seek each other out based on very specific characteristics that can be vanishingly rare in the population as a whole. In the current context, autistic individuals may dramatically increase their chances of meeting (and therefore also of forming romantic relationships) by becoming members of autism-related online discussion boards, support groups and social media pages. Such processes may dramatically increase the likelihood of autistic individuals meeting and having children (Nordsletten et al., 2016), and could therefore be an explanation (amongst many others) for why autism prevalence has risen notably in recent years.

There are several limitations to the current research that should be acknowledged. Firstly, although the sample examined is arguably more representative of the general population than most of those previously examined in this area, we did not record information relating to the presence/absence or number of offspring. Therefore, although positive assortment appears to exist within this sample, it remains unclear exactly what effects this could have on the gene pool of subsequent generations. Our study is also correlational in nature, meaning that it is not possible to assess the development of relationships over time in order to determine causal inferences (see Blossfeld & Timm, 2003). Additionally, due to COVID-19-related restrictions, only a subsample of our study participants was administered the RMET and EFT, meaning that analyses relating to these variables achieved lower statistical power than those for the AQ, EQ, and SQ-R. Although we notably still demonstrated statistically significant positive intra-couple correlations for the RMET and EFT, replication and extension of these findings will be necessary for firmer conclusions to be drawn. For instance, future studies will be required to determine whether assortative mating processes apply specifically to these variables or whether such effects are explained by within-couple similarity for IQ score.

## 5. Conclusions

The current study demonstrates small-to-moderate levels of partner similarity for a range of traits associated with autism spectrum conditions, and so implies that assortative mating may play an important role in terms of the maintenance and transmission of genes related to autistic phenotypes. In particular, we demonstrate here for the first time that systemizing is positively correlated within heterosexual partners, and that actual couples are more similar on this trait than would be expected under the assumption of a random mating pattern. These findings strongly support the assortative mating theory of autism (Baron-Cohen, 2006a, 2006b, 2007), suggest that future behavioural genetics studies should consider the influence of assortative mating when deriving heritability estimates for autism-related measures, and may be informative regarding spatial and chronological trends in autism prevalence.

## Acknowledgements

The authors would like to thank Prof. Robin Dunbar for providing useful feedback on an earlier version of the manuscript.

## Appendix Meta-analysis of the intra-couple correlation for autistic traits

We identified 14 studies in which intra-couple correlations for autistic traits were reported or could be determined from the available descriptive statistics (**Table S1**). Although not specified in our pre-registration document, we meta-analysed the available literature (k=15, n=5,770) in order to provide a more reliable effect size estimate (see **Figure S1** for forest plot). We conducted random effects meta-analyses using the R package metafor (Viechtbauer, 2010) so as to allow for the possibility of the true effect size of the correlation differing depending on moderating factors. The model revealed a statistically significant positive correlation, *z* = 0.203 (95% CI = 0.109, 0.297 [*r* = 0.200; 95% CI = 0.109, 0.289]), *p* < 0.0001, and significant heterogeneity was observed, Q (14) = 106.063, *p* < 0.0001, τ = 0.168, I^2^ = 89.18%. Removal of any one sample did not noticeably change these results (in all cases, *p* < 0.001), the funnel plot appeared reasonably symmetrical (**Figure S2**), and the trim and fill procedure (Duval & Tweedie, 2000) did not estimate the presence of missing studies.

**Table S1.**
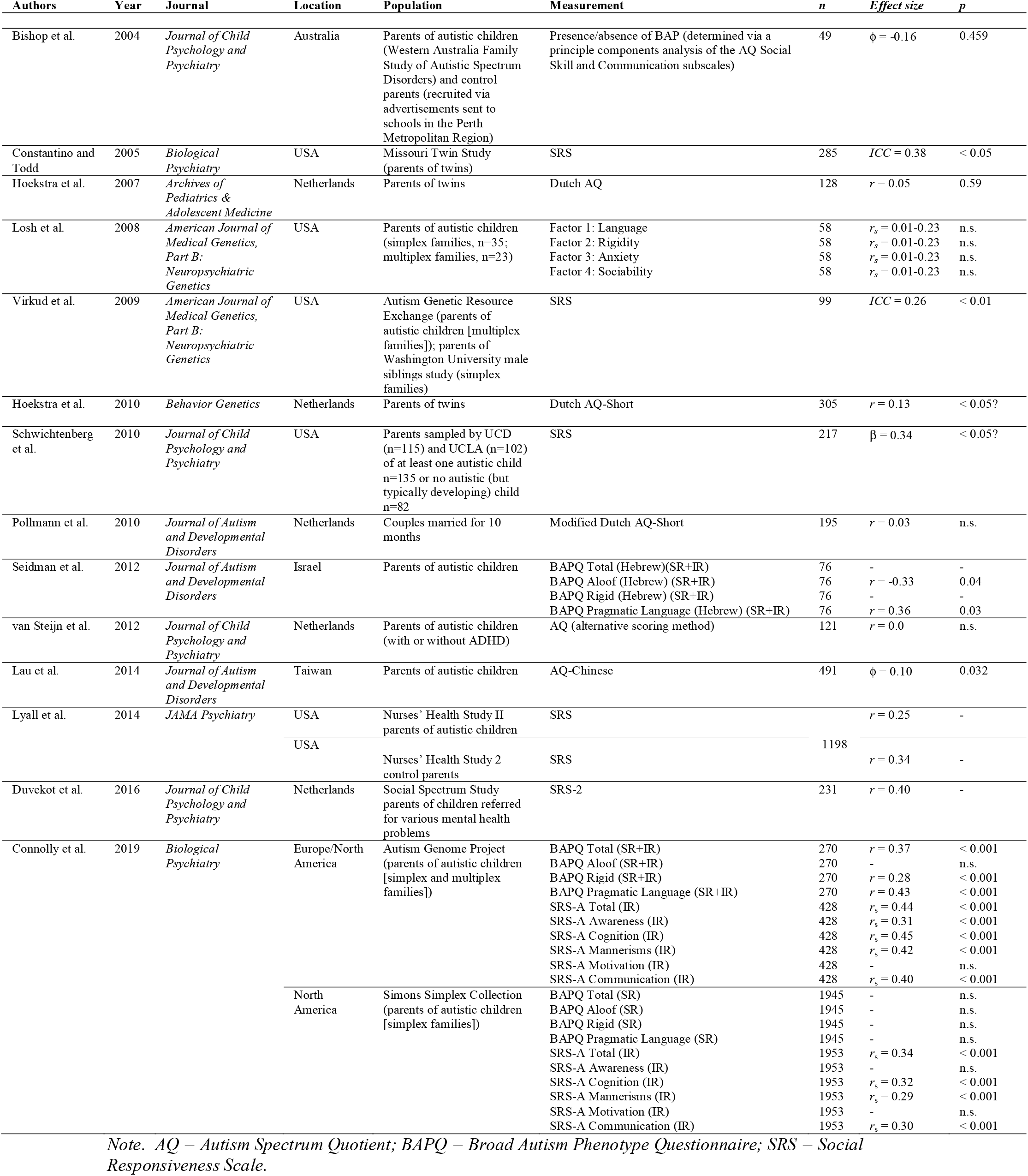
Overview of studies reporting on intra-couple correlations for autistic traits variables.

**Figure S1.**
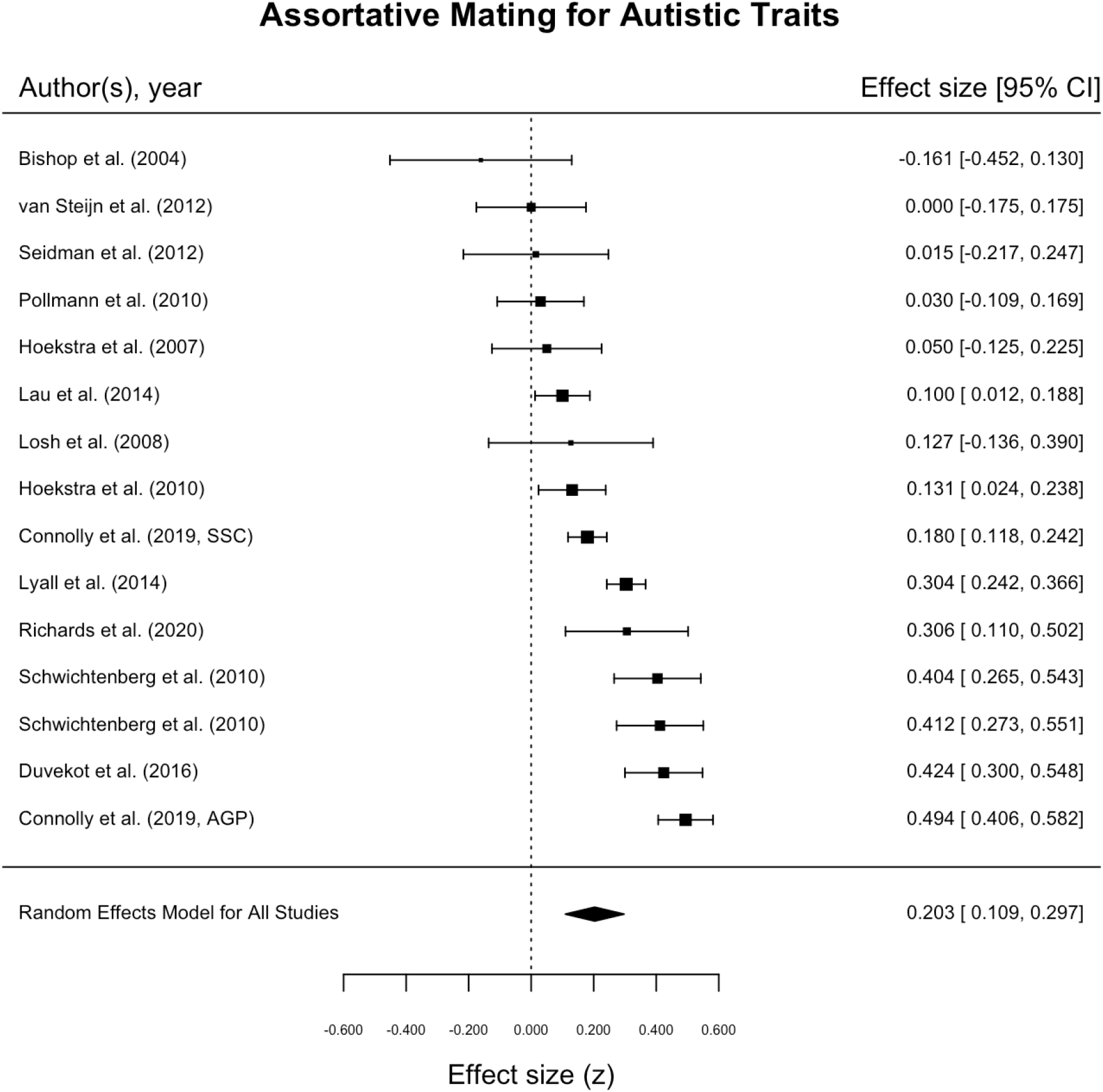
Forest plot for intra-couple correlations for autistic traits.

**Figure S2.**
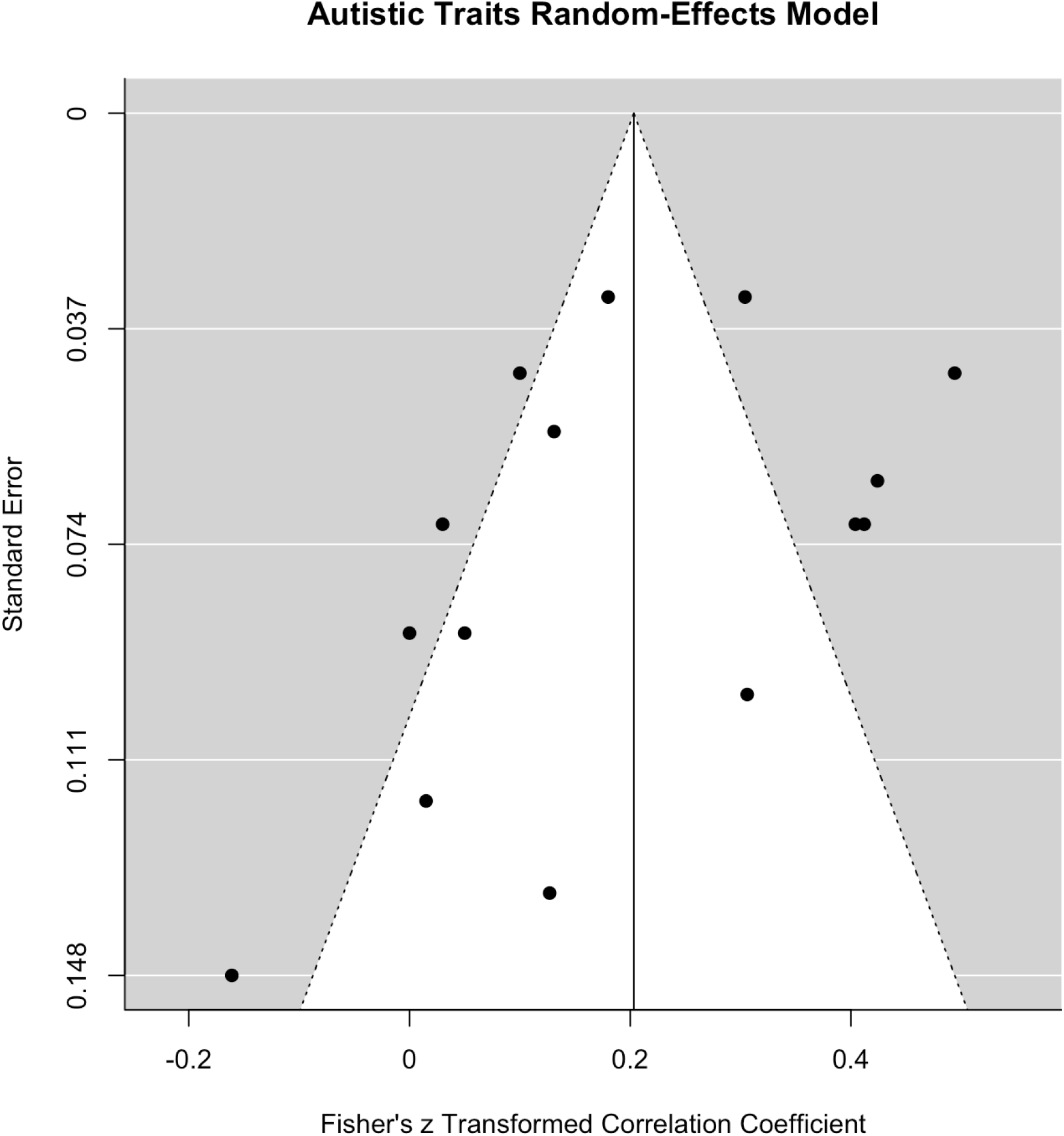
Funnel plot for studies reporting intra-couple correlations for autistic traits measures.

